## Supplemental Notes 1-5, Supplemental Tables 1 & 2, Supplemental Figures 1-14 for "Extraction of accurate cytoskeletal actin velocity distributions from noisy measurements"

### Supplementary information for: Extraction of accurate cytoskeletal actin velocity distributions from noisy measurements

**Cayla M. Miller**[a], **Elgin Korkmazhan**[b], **and Alexander R. Dunn**[a,c]✉

[a]Department of Chemical Engineering, Stanford University, Stanford, CA, USA
[b]Biophysics Program, Stanford University, Stanford, CA, USA
[c]Stanford Cardiovascular Institute, Stanford University School of Medicine, Stanford, CA, USA

#### Supplementary Note 1: Derivation of distance distribution

The analytical form for the distribution of measured distances, $d$, given a true distance, $s$, is an extension of the derivation by Churchmann et al.(1). If we assume that two points are located at (0,0) and ($s$,0), and that the measured coordinates are Gaussian distributed around the true location with variance $\sigma^2$, we can say that $x_1, y_1, y_2 \sim N(0, \sigma^2)$ and $x_2 \sim N(s, \sigma^2)$. As a result, the measured distance along x and y, $d_x$ and $d_y$, respectively, are also normally distributed.

$$d_x = x_2 - x_1 \sim N(s, 2\sigma^2)$$

$$d_y = y_2 - y_1 \sim N(0, 2\sigma^2)$$

The noncentral $\chi$ distribution describes the distribution of a random variable $z$ given by

$$z = \sqrt{\sum_i^k \left(\frac{X_i}{\sigma_i}\right)^2}$$

where $X_i$ are normally distributed random variables with means $\mu_i$ and variances $\sigma_i^2$.
Therefore we may define $z$ in terms of $d$ as a noncentral $\chi$-distributed random variable with $k$=2:

$$z = \sqrt{\left(\frac{d_x}{\sigma_{dx}}\right)^2 + \left(\frac{d_y}{\sigma_{dy}}\right)^2} = \frac{d}{\sqrt{2}\sigma}$$

Assuming $\sigma_{dx}$ and $\sigma_{dy}$ are equal, the noncentral $\chi$ distribution is

$$f_z(z, k, \lambda) = \lambda(\lambda z)^{-k/2} exp\left(-\frac{z^2 + \lambda^2}{2}\right)\left(z^k\right) I_{k/2-1}(\lambda z),$$

where $I_{k/2-1}$ is a modified Bessel function of the first kind and

$$\lambda = \sqrt{\sum_i^k \left(\frac{\mu_i}{\sigma_i}\right)^2} = \sqrt{\frac{s^2}{2\sigma^2} + \frac{0^2}{2\sigma^2}} = \frac{s}{\sqrt{2}\sigma}.$$

Dividing by $\sqrt{2}\sigma$ to determine the distribution of $d$, and substituting $k = 2$ and $\lambda = s/(\sqrt{2}\sigma)$, we find that

$$f_d(d, s, \sigma) = \frac{d}{2\sigma^2} exp\left(-\frac{1}{4\sigma^2}\left(d^2 + s^2\right)\right) I_0\left(\frac{ds}{2\sigma^2}\right)$$

#### Supplementary Note 2: Drift-diffusion and jump process models

**A. Drift-diffusion.** We modeled F-actin movement with two simple drift-diffusion models that we will collectively call the drift-diffusion model. In the first model, the particles undergo 1-D Brownian motion while drifting at a constant 1-D velocity. This results in folded Gaussian distributed distances, with location parameter $\mu$ equal to the drift velocity times the observation time and spread parameter $\sigma^2$ equal to the diffusion coefficient times observation time. The second model differs from the first only in that the particles are allowed to diffuse in 2-D while drifting in 1-D, which results in Rice-distributed distances with center $\nu$ equal to the drift velocity times the observation time and spread parameter $\sigma^2$ equal to the diffusion coefficient times observation time.

**B. Position jump.** We also modeled F-actin movement with a simple one dimensional unidirectional jump process with independent identically distributed (IID) exponential wait times between jumps and IID exponential jump distances at jumps. This function is sufficiently simple that we can obtain the solution in real space. When observed at a time interval $\tau$, the total number of jumps in the interval will be Poisson distributed with mean $\mu\tau$ where $\mu$ is the jump rate.
If $k$ non-zero jumps occur in an interval, the total distance moved in that interval will be distributed as the sum of $k$ IID exponential random variables each with some rate parameter $\beta$. This gives a Gamma distribution of distances with shape parameter $k$ and rate parameter $\beta$.
To find the probability distribution of the distance $s$ moved for a given observation time window $\tau$, we first need to sum up the probability that there is one jump of size $s$; two jumps of total size $s$; three jumps of total size $s$; and so on, giving

$$\begin{aligned}
&\sum_{k=1}^{\infty} \frac{(\mu\tau)^k}{k!} e^{-\mu\tau} \gamma(k,\beta) \\
&= \sum_{k=1}^{\infty} \frac{(\mu\tau)^k}{k!} e^{-\mu\tau} e^{-\beta s} s^{k-1} \frac{\beta^k}{(k-1)!} \\
&= \frac{1}{s} e^{-\mu\tau-\beta s} \sum_{k=1}^{\infty} \frac{(\mu\tau\beta s)^k}{k!(k-1)!} \\
&= \frac{1}{s} e^{-\mu\tau-\beta s} \sqrt{\mu\tau\beta s} I_1(2\sqrt{\mu\tau\beta s}) \\
&= e^{-\mu\tau-\beta s} \sqrt{\frac{\mu\tau\beta}{s}} I_1(2\sqrt{\mu\tau\beta s}) \qquad (1)
\end{aligned}$$

where $\gamma(k,\beta)$ is the Gamma distribution with shape parameter $k$, rate parameter $\beta$ and $I_1$ is the modified Bessel function of the first kind, of order 1. This expression is incomplete since we also need to consider the (discrete) probability of moving a distance of exactly $s = 0$, corresponding to the case of no jumps. The probability of a point moving exactly zero distance is equal to the probability that a jump does not occur in the given observation interval, such that $P(s = 0) = e^{-\mu\tau}$. Therefore, the complete normalized probability distribution for moving a distance $s \geq 0$ is defined piece-wise as

$$P(s) = \begin{cases} e^{-\mu\tau} \text{ for } s = 0 \\ e^{-\mu\tau-\beta s}\sqrt{\frac{\mu\tau\beta}{s}} I_1(2\sqrt{\mu\tau\beta s}) \text{ for } s > 0 \end{cases} \qquad (2)$$

**C. Position jump with reversals.** To model 1D position jumps with reversals, we keep the properties above (IID exponentially distributed jump distances and IID exponentially distributed wait times between jumps), but add a constant probability of reversing direction on any jump.

This model was not readily tractable in real space, and instead fitting to this model is done via iterative Monte Carlo simulation (see Methods).

#### Supplementary Note 3: Numerical approximation of Bayes Theorem (Eqn. 1)

The probability of observing a step distance, $d$, given a distribution of true distances, $f_s(s)$, is given by

$$f_d(d) = \int_{-\infty}^{\infty} f_d(d|s = S) f_s(S) dS. \qquad (3)$$

While this equation may be expressed analytically for some forms of $f_s(s)$, we fit to a numerical approximation of this equation for consistency and ease of application to various functional forms. For our case, where we fit to distances, no values are less than zero, and the limits become zero to infinity. We approximate this integral as a Riemann sum over the interval from 0 to 1000 nm, with 100 linearly-spaced bin edges (bin width = 10 nm). For each bin, both $f_s(S)$ and $f_d(d|s = S)$ are calculated at both bin edges, averaged, and multiplied by the bin width. These are summed to give $f_d(d)$.

For the jump process as described in Appendix 2, Bayes rule is not defined exactly as above due to the discrete discontinuity at $s = 0$. Instead, because the probability is defined piece-wise (Eqn. 2), Bayes theorem is rewritten in this case to give

$$f_d(d) = f_d(d|s = 0) f_s(s = 0) + \int_{0+}^{\infty} f_d(d|s = S) f_s(S) dS. \qquad (4)$$

We approximate the integral here by approaching zero from the right, using 100 linearly-spaced bin edges from 0.0001 to 1000 nm.

#### Supplementary Note 4: Estimation of F-actin lifetimes

In order to estimate the lifetimes of our measured F-actin populations, we analyzed the lifetimes of our tracked F-actin puncta. We begin with the fixed-cell puncta as a baseline of non-turnover based lifetimes (i.e. lifetimes arising from SiR-actin binding kinetics, bleaching, etc.). Lifetimes in fixed cells were well fit by a biexponential with rates $r_1 = 0.06/s$ and $r_2 = 0.33/s$ (Supplementary Fig. 11, left). While typical reversible binding is expected to yield only a single exponential, these two rates may arise from SiR-actin (based on the actin binding moiety from jasplakinolide) binding to F-actin in at least two modes, potentially dependent on F-actin nucleotide state(2).

We then fit the puncta lifetimes measured in live cells to a biexponential. Because these rates are expected to arise from the combined rates of bleaching, SiR-actin unbinding and F-actin turnover, the two rates are given by $r_1 + r_{to}$ and $r_2 + r_{to}$, where $r_1$ and $r_2$ are the rates from the fixed population fits, and $r_{to}$ is the turnover rate of speckles in a given population. These fits yielded F-actin turnover rates of 0.06/s for cortical F-actin and 0.03/s for stress fiber F-actin, giving lifetimes of 18 and 34 s, respectively (Supplementary Fig. 11, middle and right). These turnover rates represent turnover both from depolymerization and movement out of the TIRF excitation field, and are comparable with previous FRAP measurements(3–6).

#### Supplementary Note 5: Robustness testing

We analyzed and compared subsets of the data to probe the possible effects of various forms of error from particle tracking. First, to test whether high speckle density in certain regions of the cell might lead to tracking errors, we compared measured displacements from high density images to measured displacements from low density images. Because the SiR-actin signal bleached significantly during the acquisition, displacements measured during the first half of the movie were measured at higher speckle densities than those measured in the second half. When we compared these displacements (at the 10 s timescale, where displacements became more easily distinguishable from localization error), the distributions were in good agreement with one another (Supplementary Fig. 12). These data suggest that the particle tracking approach used here is robust to variations in fluorophore density.

As an independent test of whether high speckle density regions have greater tracking errors (e.g. false linkages from overlapping fluorophores), we compared measured displacements of the brightest 25%, the dimmest 25% and the remaining 50% of tracks, where brightness was determined from the average amplitude of the best fit Gaussian during subpixel localization. These displacements were greatest for the dimmest tracks and least for the brightest tracks, likely because dim fluorophores were more poorly localized than bright ones. We tested the robustness of our fits by fitting each of these brightness populations to a Weibull distribution, though without updating our noise model to more accurately reflect the localization error of those tracks. Best fits to these data yielded shape parameters between 1 and 1.5, consistent with motion being more Poisson-like than diffusive, and similar shape parameters for all three brightness populations (Supplementary Fig. 13).

Last, we compared fits from tracks of varying lifetimes. For any given time between a pair of localizations (e.g. 10 s apart), measured displacements may come from tracks of varying lengths: those that last only the given time delay, and those that last much longer. When fit to a Weibull, we found that for a given observation timescale, tracks of varying durations yielded similar shape parameters, but with some variation in scale parameter, with the shortest duration tracks yielding the greatest scale parameters (Supplementary Fig. 14). It is possible that this observation reflects greater mobility and faster turnover for shorter filaments, as might be expected if filaments are restrained by crosslinks that are randomly and uniformly distributed along their lengths. However, it is also possible that this result may reflect experimental factors, for example the fact that brighter fluorophores were more likely to be successfully tracked over a long period of time than dim ones. To further probe this observation, we fit both stress fiber and cortical populations to the 1D jump model with reversals, including either all tracks or only long (> 14 s) tracks, and found that while there were changes in the best fit parameters, overall trends held (i.e. the cortical population was best fit by shorter time between jumps and lower reversal probability than tracks from stress fibers, Supplementary Table 2). Thus, despite some possible differences, the movement of both short- and long-duration tracks was qualitatively similar, and well-described by a jump model.

**Table S1.** Representative log10(likelihood-ratios) for several two-parameter model comparisons, from one experimental replicate. Abbreviations are as follows: Weibull: Wbl, folded Gaussian: Fgaus. Jump represents the 1D jump model, without reversals.

| Population | Wbl vs. Fgaus | | Jump vs. Rice | | Wbl vs. Jump | | Jump vs. Fgaus | |
|---|---|---|---|---|---|---|---|---|
| timescale | 20 s | 40 s | 20 s | 40 s | 20 s | 40 s | 20 s | 40 s |
| SFs | 76 | 1.9 | 377 | 53 | 14 | 0.9 | 62 | 1.0 |
| cortical | 47 | 6.5 | 343 | 51 | 17 | 4.0 | 30 | 2.5 |
| adhesions | 11 | 5.3 | 71 | 2.5 | 4.1 | 4.0 | 6.7 | 1.2 |
| adhesions$\wedge$SFs | 3.8 | 1.9 | 45 | 11 | 2.1 | 0.9 | 1.7 | 0.9 |

**Table S2.** Best-fit fit parameters for all tracks vs. only long tracks, fit to a 1D jump model with reversals. Data fit is from both stress fiber and cortical populations, from one experimental replicate.

| | All tracks | | Tracks > 14 s | |
|---|---|---|---|---|
| Fit parameter | Stress fibers | Cortical | Stress fibers | Cortical |
| average time between jumps (s) | 6.9 | 3.6 | 8.9 | 5.6 |
| average jump distance (nm) | 54.5 | 61.5 | 49.4 | 61.3 |
| reversal probability | 0.26 | 0.37 | 0.03 | 0.19 |

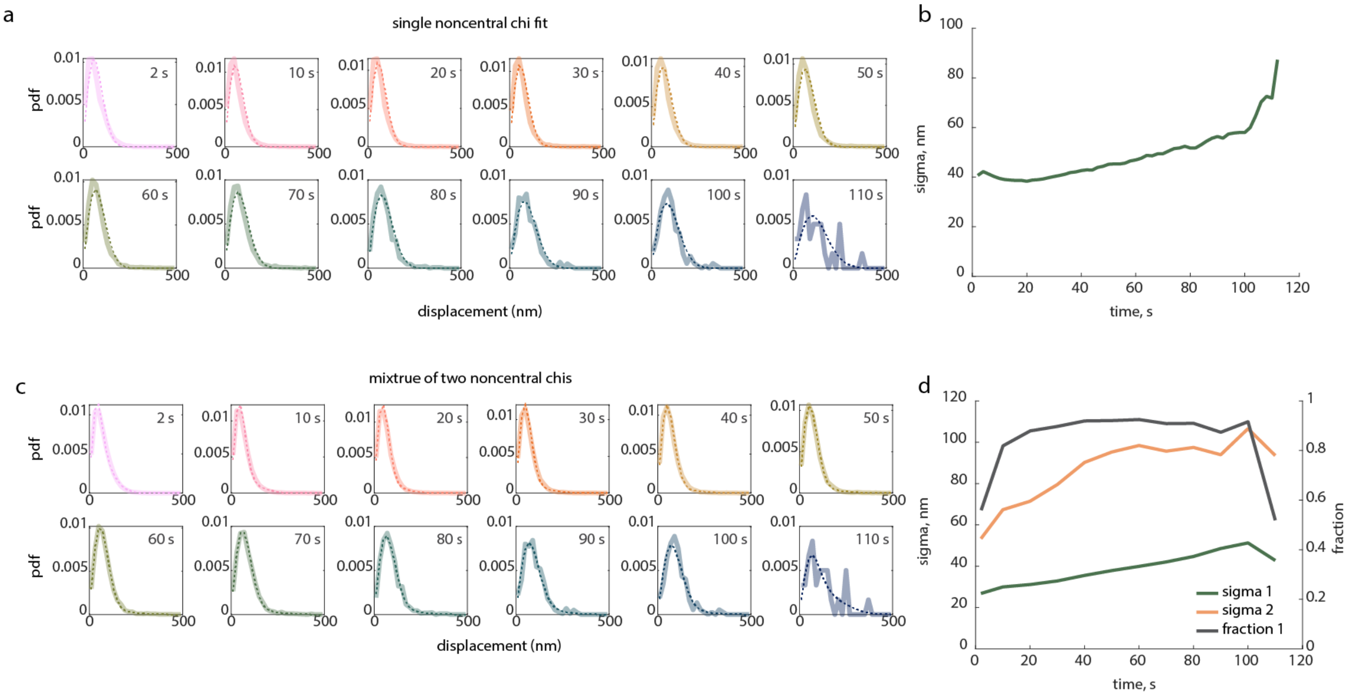


**Fig. S1.** a,c) Distance distributions for fixed HFFs, with varying time intervals between frames (time interval, in seconds, indicated in the top right corner of each). The best fit distributions to a single noncentral chi (a) or a mixture of two noncentral chi distributions (c) with s=0 are shown in dashed lines. b,d) The best fit parameters for each timescale. We take the shortest timescale fit as the best estimate of our localization error, as it represents our most static sample, and use this in our downstream workflow. n = 10 cells.

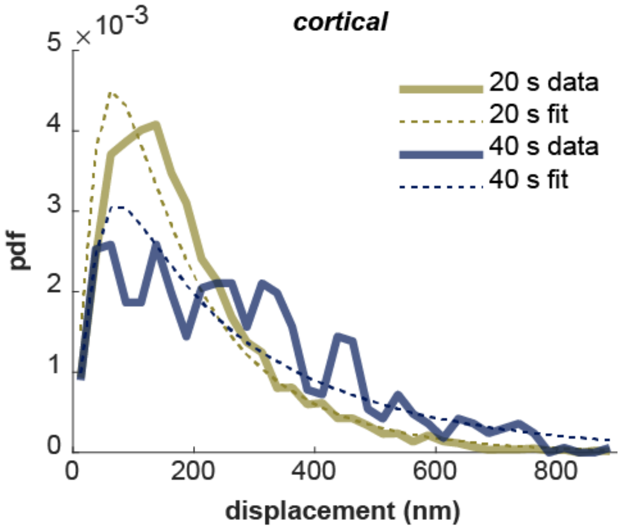


**Fig. S2.** Exponential fits to the cortical population in HFF, at 20 and 40 s timescales show poor fits to the data. n = 8 cells.

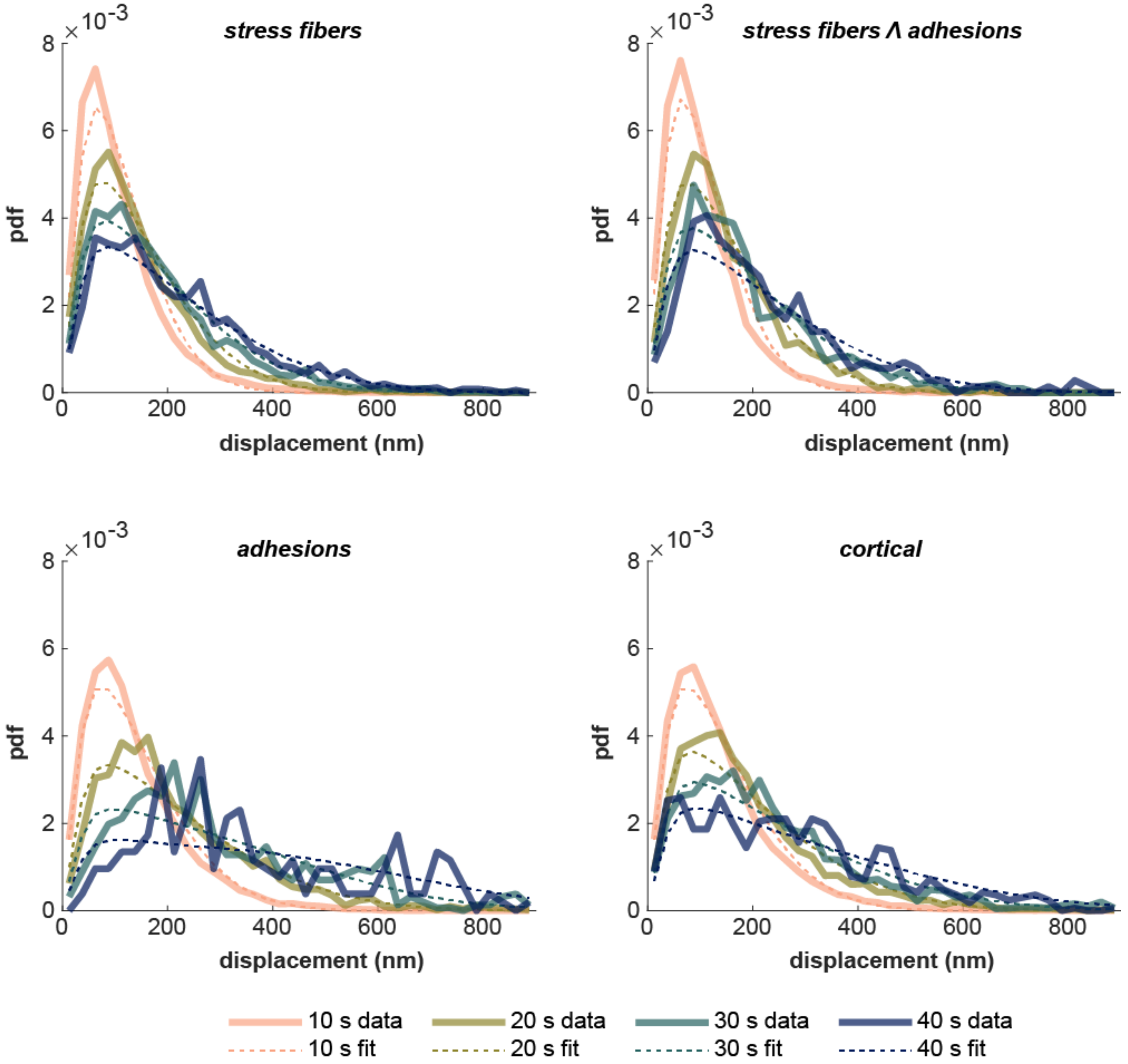


**Fig. S3.** Folded Gaussian fits to all four populations over varying timescales (10, 20, 30, and 40 s). The folded Gaussian is less heavy-tailed than the measured distribution, and tends to either underestimate the tails (most noticeable at the 10 s timescale) or miss the peak in order to fit the tails (e.g. the adhesion and cortical fits at longer timescales). n = 8 cells.

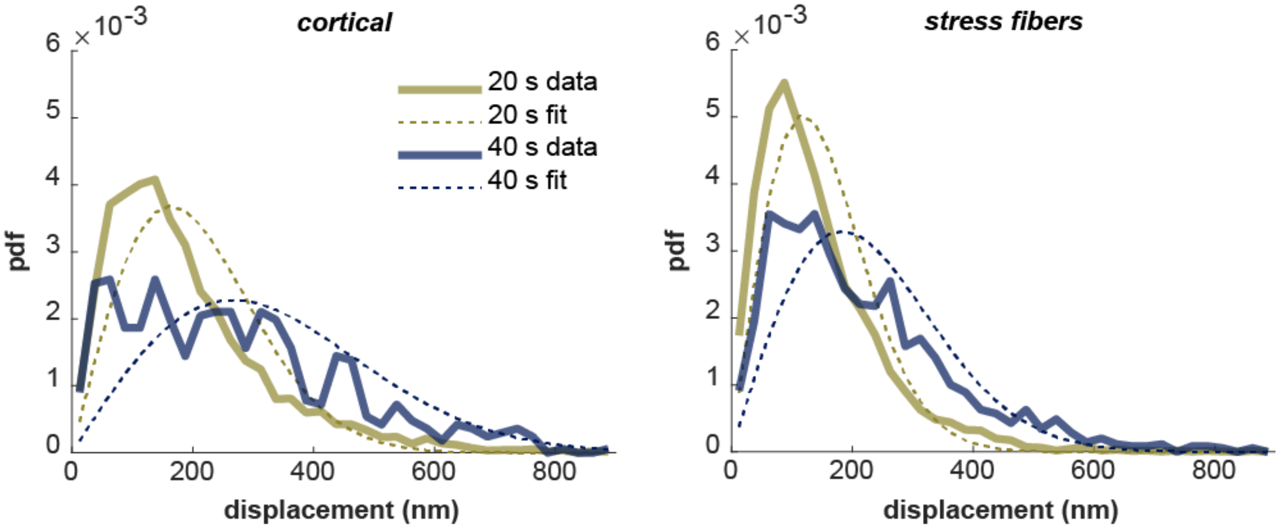


**Fig. S4.** Rayleigh fits to both stress fiber and cortical populations in HFFs at the 20 s and 40 s timescales show poor fits to the data. n = 8 cells.

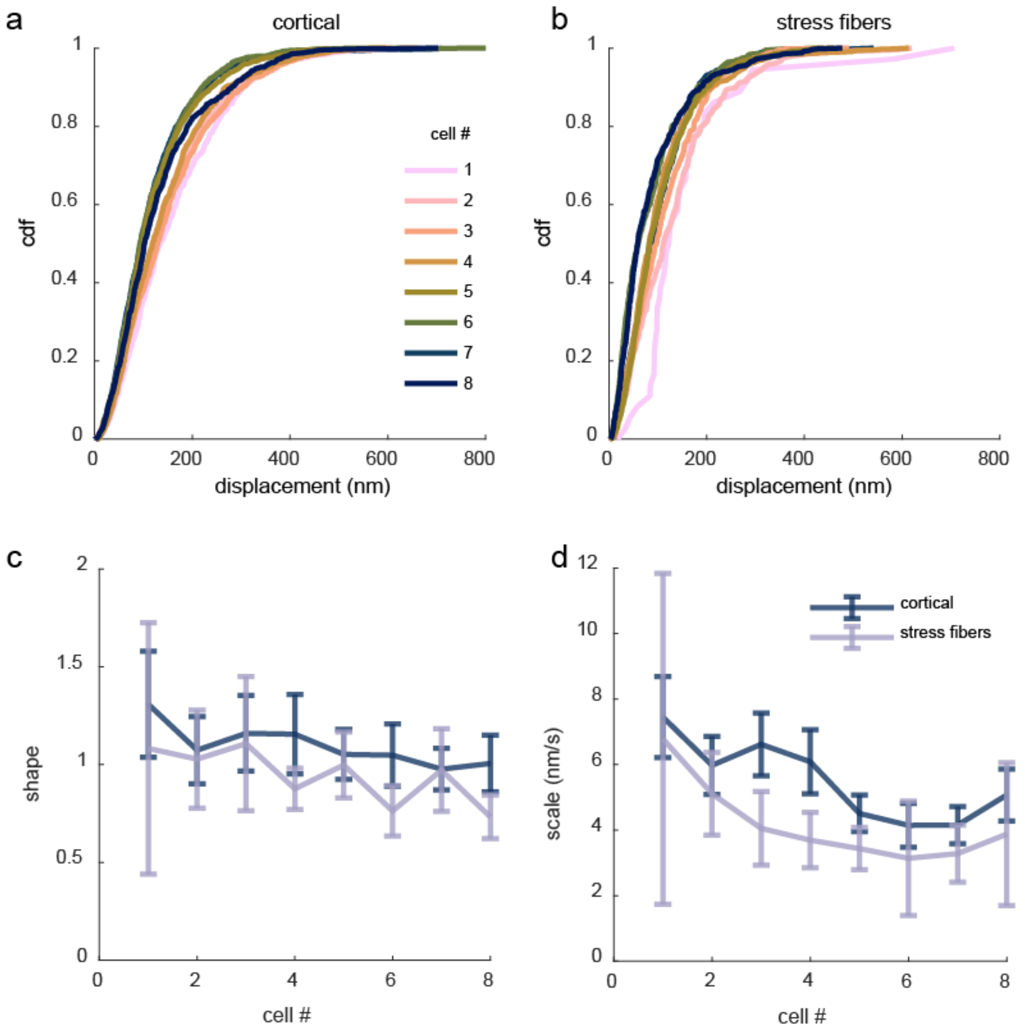


**Fig. S5.** a,b) Cumulative distribution functions of raw (uncorrected) displacement distributions for cortical (a) and stress fiber (b) populations in HFFs for eight separate cells. All cdfs are for a 20 s time interval. c,d) Best fit parameters for a Weibull distribution fit to the displacement distributions in a and b. Error bars show the 95% confidence interval for each parameter, determined by normal approximation (see Materials and Methods, Model Fitting). n = 8 cells.

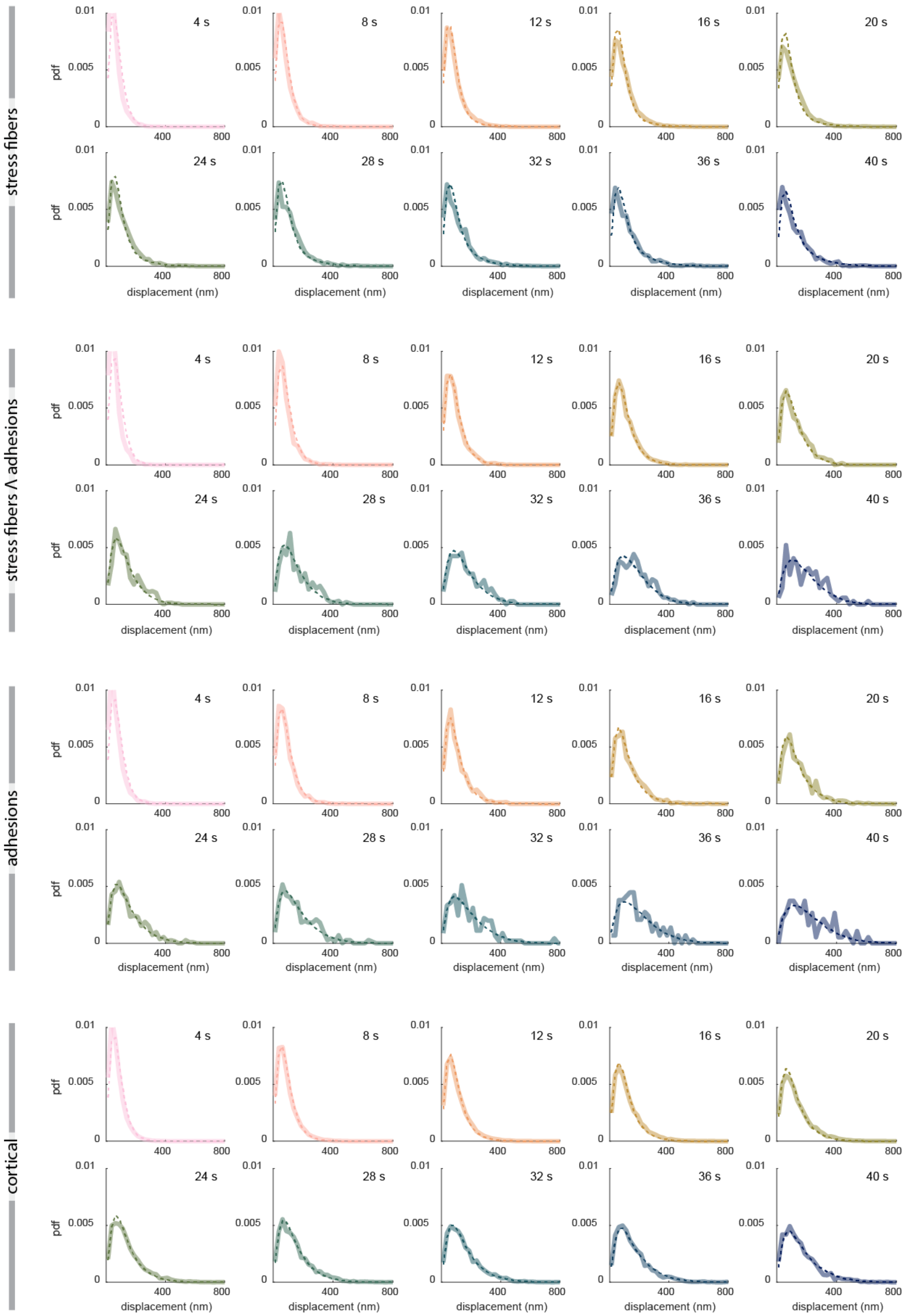


**Fig. S6.** Example goodness of fit for a fit of the 1D position jump process with reversals to the four actin populations in HFFs, representative of 1 experiment with n=9 cells. All timescales are fit simultaneously by one set of fit parameters. The measured distributions are shown in thick, semitransparent lines, and the best fits are shown in dashed lines.

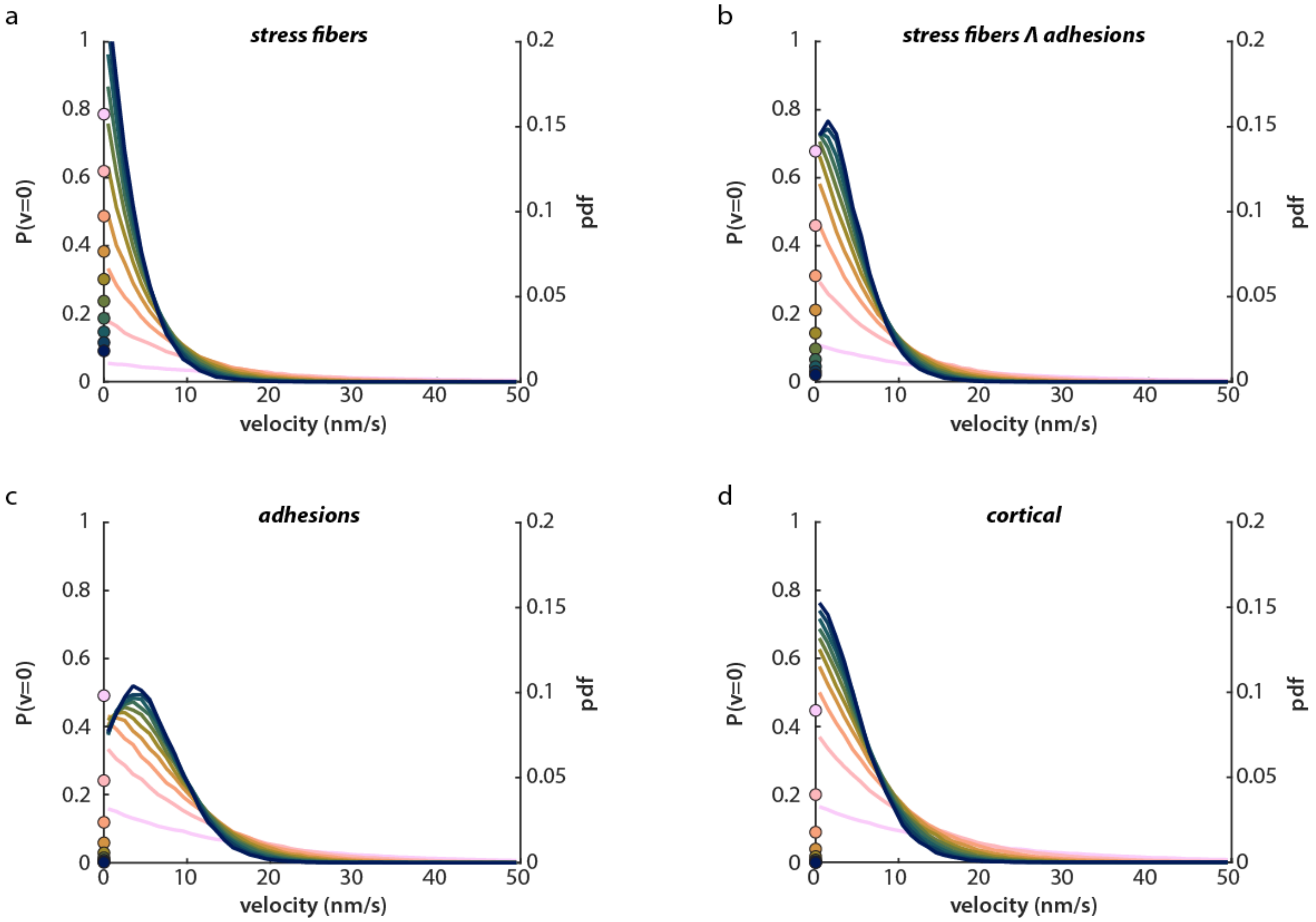


**Fig. S7.** Inferred velocity distributions from the 1-D jump model with reversals for all four F-actin populations in HFFs. The distributions for each cellular population are generated from the average best fit parameters (Fig 4g) across all HFF datasets (n = 32 total cells from 4 experimental replicates). In each case, the discrete probability of zero velocity is shown on the left y-axis, while the right y-axis gives the pdf values for the continuous pdf at velocities greater than zero.

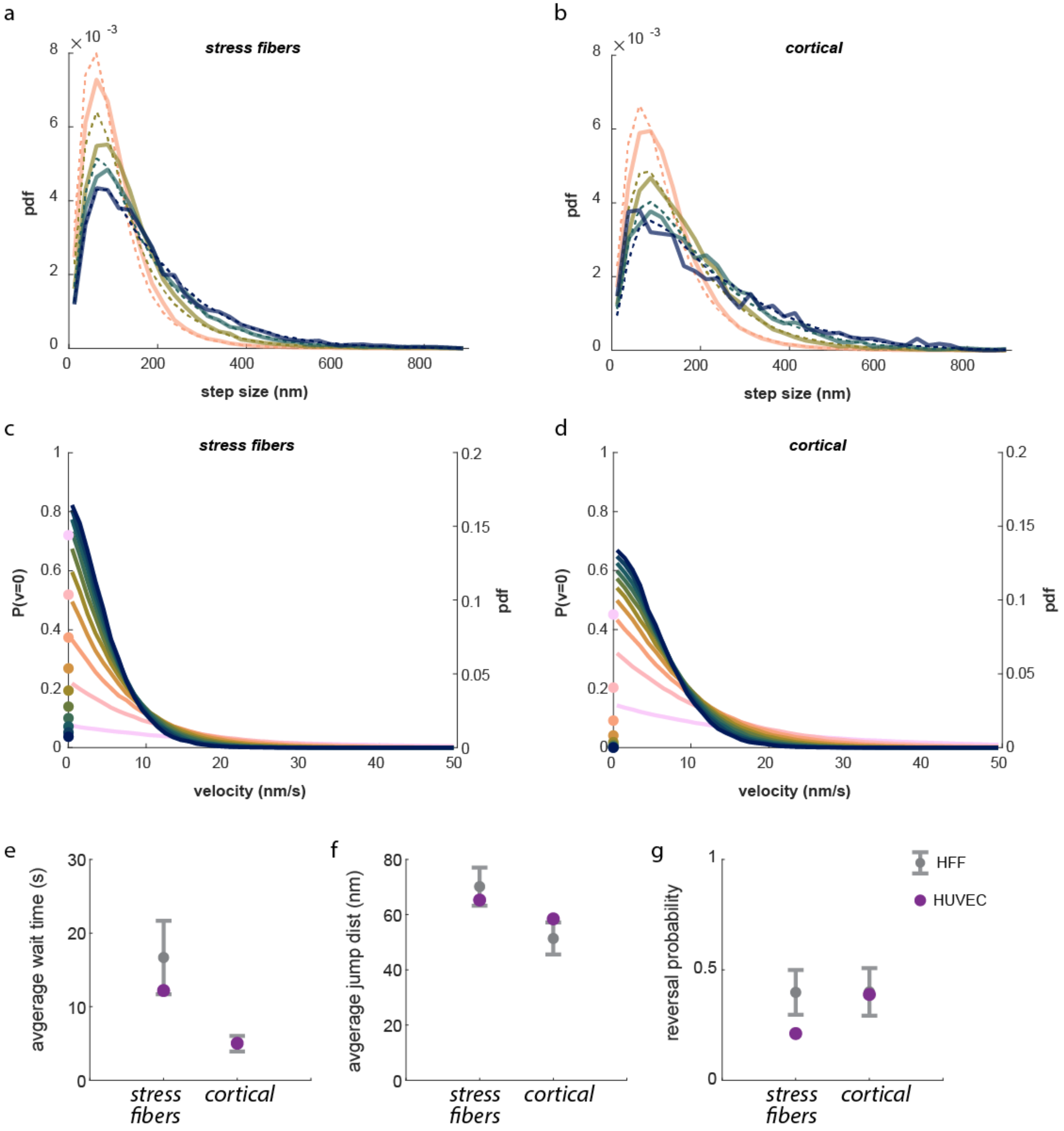


**Fig. S8.** a,b) Measured and fit probability density functions for stress fibers (a) and cortical actin (b) in HUVECs for varying timescales (10, 20, 30, 40 s; solid lines), and the corresponding best fit for each (dashed lines) to the 1D position jump with reversals. c,d) The inferred velocity probability distributions for the fits to the distributions in (a) and (b). e,f,g) The best fit parameters for each fit (shown in a-d) for HUVECs (purple), as compared to those for HFFs (gray) shows that HUVECs share similar fit parameters with HFFs. n = 8 cells.

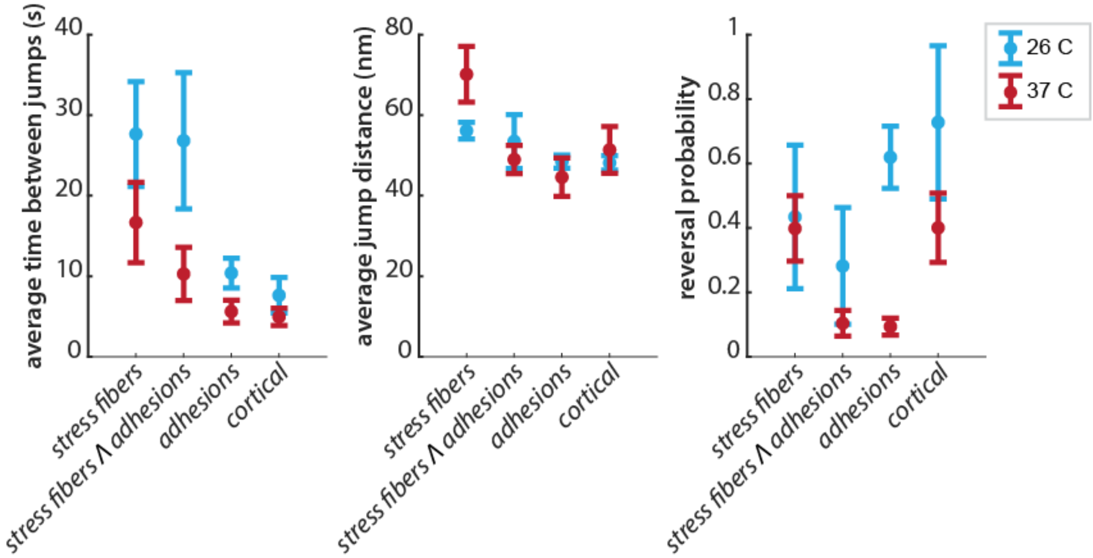


**Fig. S9.** Temperature dependence of best fit parameters of the 1D position jump with reversals in HFFs. Measurements were made at both 37C (red) and 26C (blue). Points represent the average best fit parameters across 3-4 experiments (26 C, 3 experiments with n = 29 total cells; 37 C, 4 experiments with n = 32 total cells). Error bars represent the standard error on the mean from independent experiments.

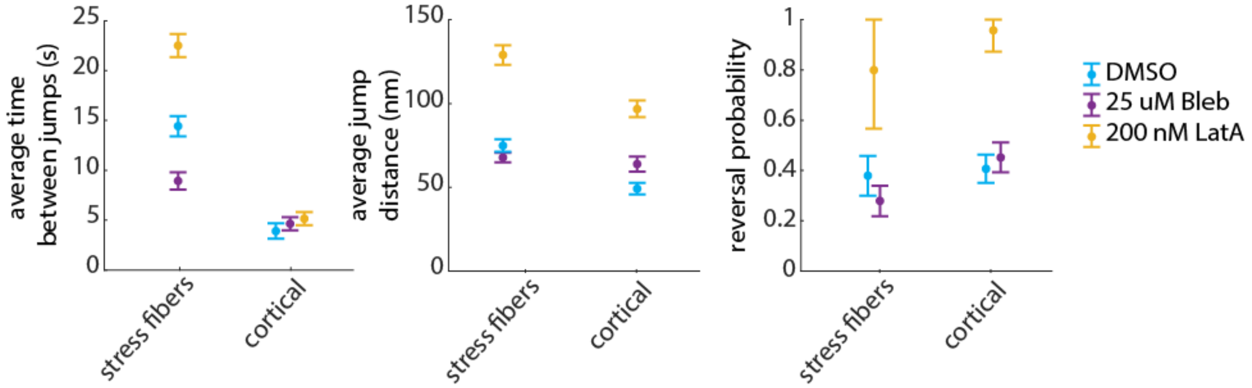


**Fig. S10.** Best fit parameters of the 1D position jump with reversals to actin tracks in HFFs treated with 0.1% DMSO (cyan, n = 9 cells), 25 $\mu$M Blebistatin (purple, n = 11 cells), or 200 nM Latrunculin A (yellow, n = 11 cells). Errorbars represent the 95% confidence intervals of fits.

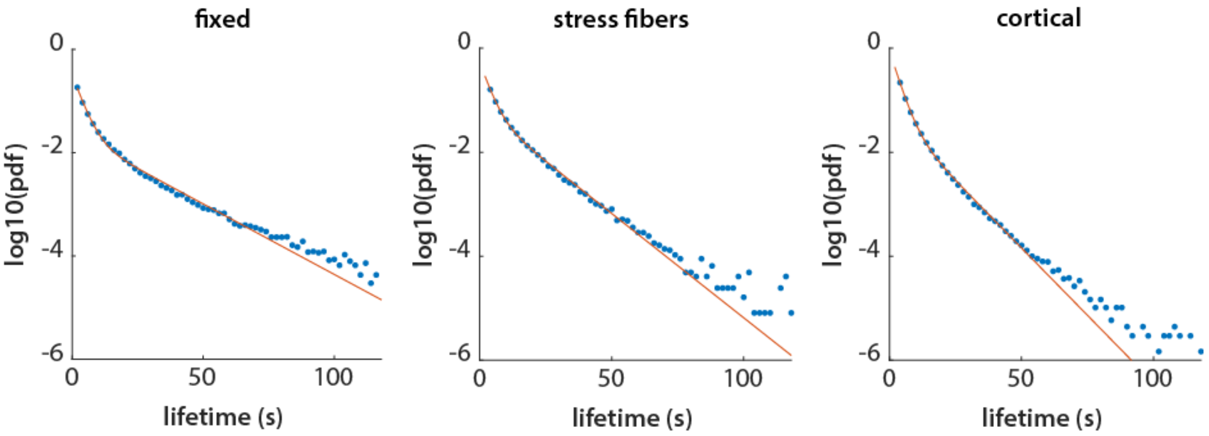


**Fig. S11.** Measured lifetimes of SiR-actin speckles (circles) in fixed cells (left, n = 10 cells), and live cell stress fiber (middle) and cortical (right) populations (n = 9 cells), as well as biexponential fits (solid lines) as described in Supplementary Note 4.

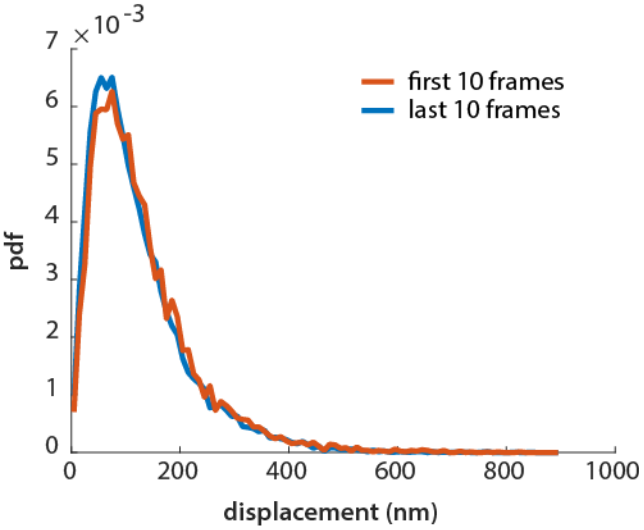


**Fig. S12.** Measured displacements in live cells (all subcellular populations) over a 10 s time interval, measured either during the first 20 s (10 frames) of a movie, or from the last 20 s of the movie. n = 9 cells.

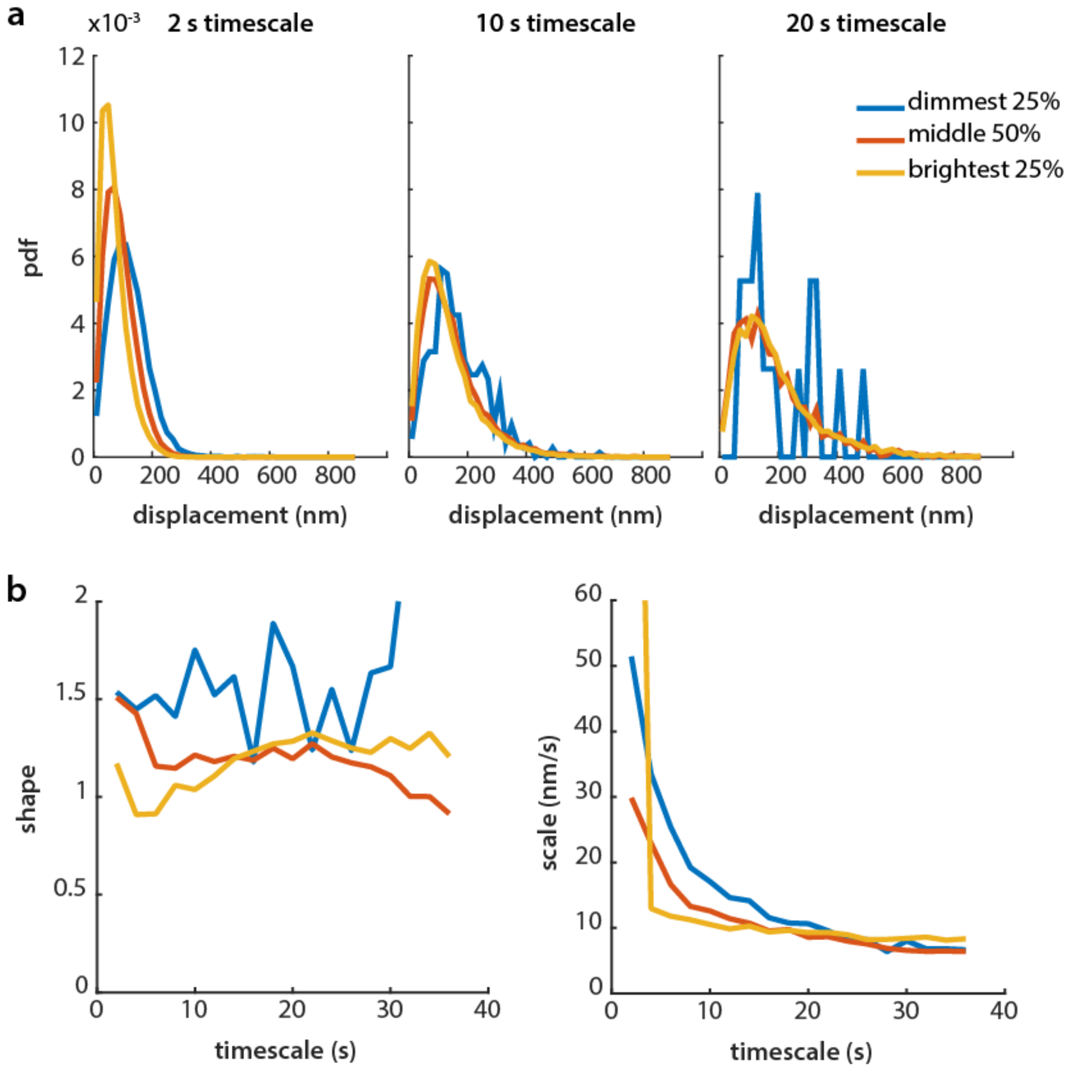


**Fig. S13.** Displacements of tracks with varying punctum brightness and Weibull fits to the same. a) Measured displacements at three timescales (2 s, left; 10 s, center; and 20 s, right) for the dimmest 25% (blue), the brightest 25% (yellow), and the remaining 50% (orange) of tracked puncta. b) Best-fit Weibull parameters to the same populations, across timescales from 2 s to 36 s. These data represent the cortical population from one experimental replicate, n = 9 cells.

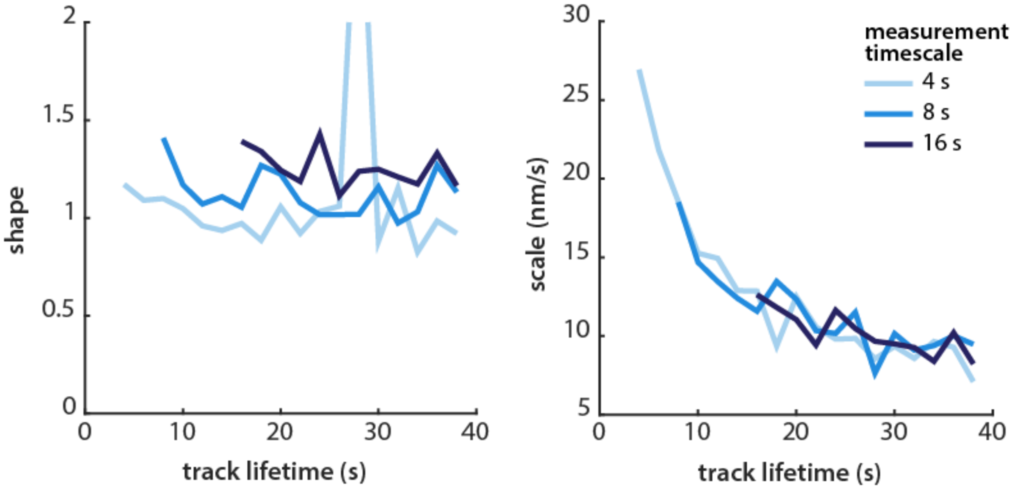


**Fig. S14.** Best-fit Weibull parameters for tracks of varying durations, measured at three time intervals: 4 s (lightest blue), 8 s, and 16 s (darkest blue). These data are drawn from the cortical population from one experimental replicate, n = 9 cells.

**Video S1. An HFF labelled with a low dose of SiR-actin.** Left: the whole cell is imaged every 2 s for 120 s total. The scale bar is 10 $\mu$m. Right: the two boxed regions of the cell on the left, magified 6x. The scale bar is 1 $\mu$m.

**Video S2. Speckle tracking by QFSM and subpixel localization** An HFF labeled with SiR-actin, imaged every 2 s for 120 s total, overlaid with tracked speckles from QFSM (circles). Localizations which pass subpixel localization are shown in green; those that fail are shown in magenta (see Materials and Methods, Actin tracking analysis for details). Video has been denoised using noise2void.
